# A Highly Scalable and Rapidly Deployable RNA Extraction-Free COVID-19 Assay by Quantitative Sanger Sequencing

**DOI:** 10.1101/2020.04.07.029199

**Authors:** Devon Chandler-Brown, Anna M. Bueno, Oguzhan Atay, David S. Tsao

## Abstract

There is currently an urgent unmet need to increase coronavirus disease 2019 (COVID-19) testing capability to effectively respond to the COVID-19 pandemic. However, the current shortage in RNA extraction reagents as well as limitations in qPCR protocols have resulted in bottlenecks in testing capacity. Herein, we describe a novel molecular diagnostic for COVID-19 based on Sanger sequencing. This assay uses the addition of a frame-shifted spike-in, a modified PCR master mix, and custom Sanger sequencing data analysis to detect and quantify SARS-CoV-2 RNA at a limit of detection comparable to existing qPCR-based assays, at 10-20 genome copy equivalents. Crucially, our assay was able to detect SARS-CoV-2 RNA from viral particles suspended in transport media that was directly added to the PCR master mix, suggesting that RNA extraction can be skipped entirely without any degradation of test performance. Since Sanger sequencing instruments are widespread in clinical laboratories and commonly have built-in liquid handling automation to support up to 3840 samples per instrument per day, the widespread adoption of qSanger COVID-19 diagnostics can unlock more than 1,000,000 tests per day in the US.

## Introduction

Coronavirus disease 2019 (COVID-19) caused by severe acute respiratory syndrome coronavirus 2 (SARS-CoV-2; COVID-19) is a public health emergency of international concern. This virus first emerged in Wuhan, China in December 2019 [1], and spread to over 160 countries [2]. As of April 6, 2020, more than 1.3 million individuals are infected and over 70,000 deaths are reported [3]. One study found that undocumented infections were the infection source for 79% of documented cases [4]. Evidence from other reports suggests that some asymptomatic or minimally symptomatic patients have high levels of viral shedding [5,6]. The substantial undocumented infection, along with the nonspecific clinical features and uncertainties in transmissibility and virulence of SARS-CoV-2, have presented significant challenges in containment of this virus. Until an effective vaccine or treatment is available, rapid, accurate, and widely available diagnostic tests not only for symptomatic patients but also for those who have come in contact with any positive cases are critical to curb the spread of the virus. However, such tests have been in limited supply in the US and many other countries in the world.

The backbone of COVID-19 diagnosis has been quantitative polymerase chain reaction (qPCR) tests. The limited number of qPCR machines and the limitations of qPCR protocols, which do not have safe stopping points or a readily automatable process, have hampered large-scale testing. Furthermore, the sensitivity of qPCR requires specific RNA extraction reagents that are currently in short supply. Other diagnostics for COVID-19 include blood-based immunoassays, which are often less accurate than molecular assays and indirectly detects immune response to COVID-19 only after symptoms appear, and CRISPR-based assays that are still in development [7].

In this report, we describe a molecular diagnostic method for COVID-19 based on quantitative Sanger (qSanger), a technique for reconstructing quantitative data from Sanger sequencing chromatograms that we originally developed and validated for non-invasive prenatal testing (NIPT) of cell-free DNA (cfDNA) [8]. The workflow of qSanger-based COVID-19 is highly similar to traditional Sanger sequencing of reverse transcription (RT)-PCR amplicons (Fig. 1), but it includes the addition of a frame-shifted synthetic COVID-19 spike-in DNA in the reaction master mix. qSanger COVID-19 is designed to support one-step reverse-transcription (RT)-PCR directly from viral transport media (VTM) of specimens, without an RNA purification step (Fig. 1A). The amplification products are then purified and Sanger sequenced by automated capillary electrophoresis. Synthetic DNA included in RT-PCR master mix prior to PCR amplification serves as an internal control that enables specimens to be readily identified as either positive or negative for COVID-19 (Fig. 1B-C). Quantitative analysis of the Sanger sequence chromatogram gives qSanger an extremely high sensitivity and specificity for all positive results with a limit of detection of 10-20 genome copy equivalents (GCE), equivalent to gold-standard qPCR methods. Furthermore, the presence of a spike-in as an intra-sample control in the qSanger assay allows for easy interpretation of results and determination of sources of error (e.g. extraction or amplification or sequencing failure), and also allows population-level analyses such as mutational analysis and contact tracing. In addition, the ratio of the amplitudes of corresponding bases between the endogenous and spike-in sequences at offset positions reflects the ratio of the molecular abundances of the two sequences. Computationally combining the amplitude ratios of multiple corresponding bases can then be used to estimate the viral load over a 400-fold dynamic range with Poisson-limited coefficient of variation.

**Figure 1.**
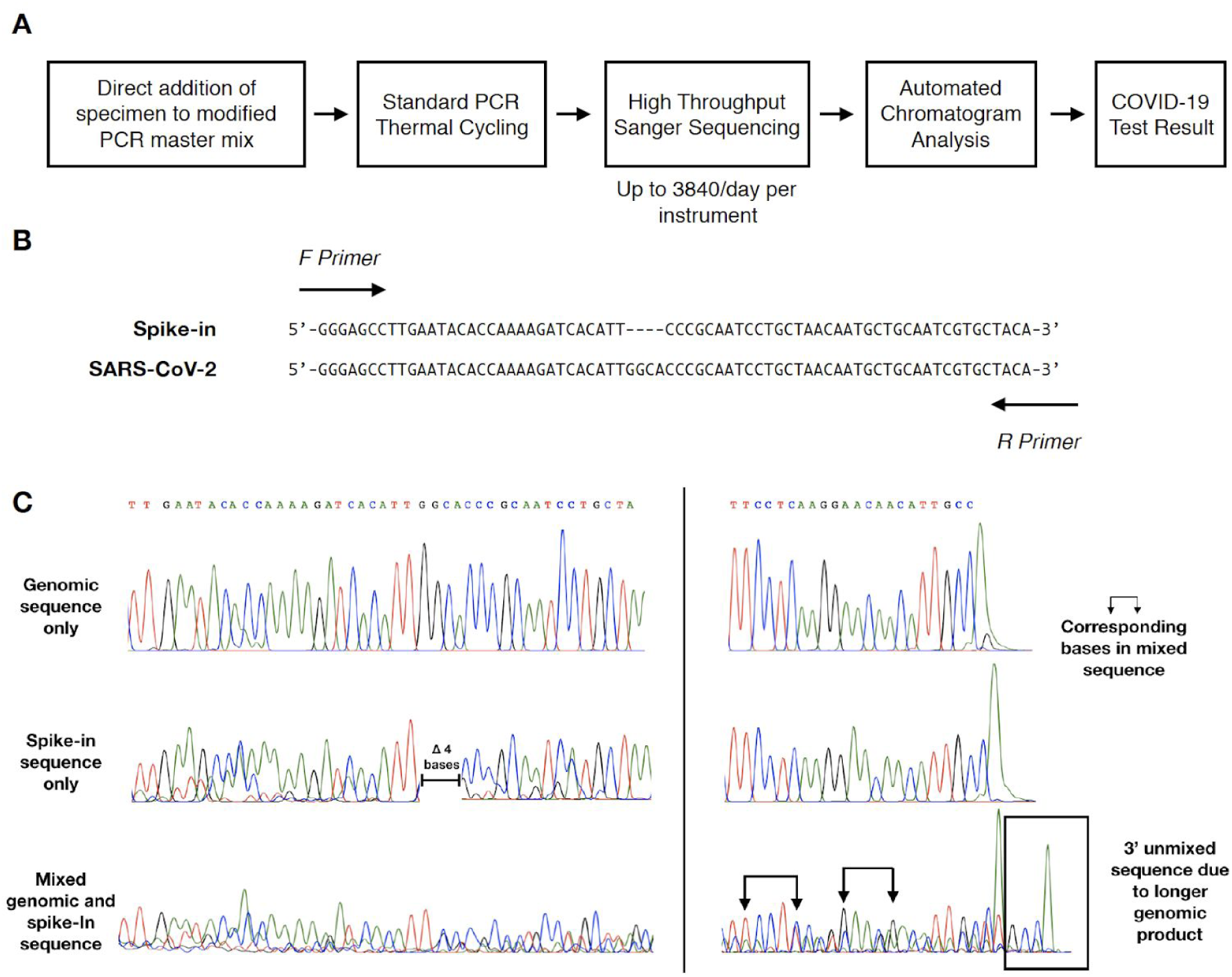
A schematic illustration of qSanger COVID-19 assay. (**A**) Specimen processing workflow. Reverse-transcription (RT) and PCR amplification of a SARS-CoV-2 target region is accomplished by directly addition of Viral Transport Media (VTM) to a one-step RT-PCR master mix containing ~200 copies of synthetic spike-in DNA. The SARS-CoV-2 target region and spike-in DNA are co-amplified on a standard thermal cycler, and the amplification products are Sanger sequenced. Custom data analysis of the resulting chromatogram is then used to determine whether the specimen is COVID-19 negative or positive. (**B**) Synthetic spike-in is designed with sequence homology to the SARS-CoV-2 target so that it co-amplifies with the SARS-CoV-2 target. A 4-base pair (bp) deletion in the spike-in design enables quantification of relative abundances of spike-in and SARS-CoV-2 DNA from a Sanger sequencing chromatogram. The depicted forward and reverse primers are not to scale. (**C**) Representative Sanger sequencing traces showing pure genomic sequence (top), pure spike-in sequence (middle), and sequencing from a mixture of genomic and spike-in sequences (bottom). The spike-in used has a 4-bp offset compared to wild type (wt), which means that when two sequences are present, the signal from each sequence can be used to estimate their relative abundances (see arrows for examples of paired bases).

## Results

### qSanger COVID-19 Limit of Detection is comparable to RT-qPCR

As an initial demonstration of qualitative detection of COVID-19 by qSanger, we designed PCR primers and synthetic DNA spike-in to target SARS-CoV-2 N protein (Fig. 1B). A one-step RT-PCR mix (NEB) containing both was used to perform reverse-transcription of SARS-CoV-2 RNA and subsequent PCR amplification in one pot. In each RT-PCR, we added either nuclease-free water as a no-template control (NTC) or 100-5000 GCE of synthetic SARS-CoV-2 RNA (Twist Biosciences). All reactions also contained ~200 GCE of spike-in in the RT-PCR master mix. After RT-PCR and Sanger sequencing, we observed a qualitatively clean chromatogram for the spike-in sequence for the NTC condition in which no SARS-CoV-2 RNA was added (Fig. 2A). At the 100 GCE level, the Sanger chromatogram showed clear mixed bases corresponding to approximately equal abundance of spike-in DNA and SARS-CoV-2 RNA. At 5000 GCE SARS-CoV-2 RNA, the chromatogram exhibited a relatively pure trace for the SARS-CoV-2 target sequence, suggesting that the SARS-CoV-2 signal overwhelmed the spike-in signal when it was present at 50-fold greater abundance.

**Figure 2.**
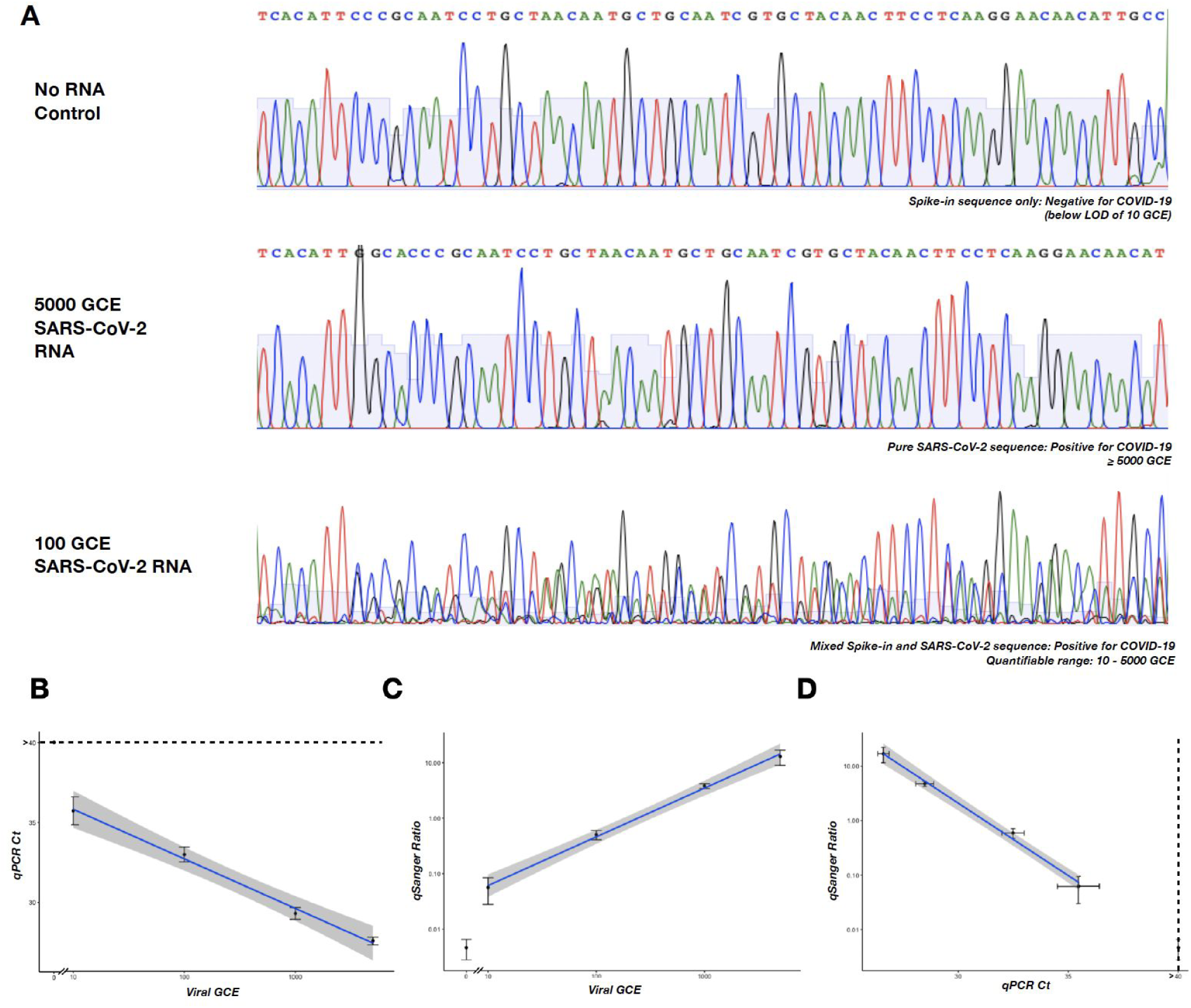
qSanger is a quantitative approach to measuring viral abundance in a sample. (**A**) Data analysis and interpretation in a qSanger COVID-19 assay. Representative Sanger sequencing chromatograms are shown for amplified products of spike-in DNA only, SarS-CoV-2 RNA only, or a mixture of the two. Since the spike-in is an internal control for amplification and sequencing, observing the spike-in signal only indicates that SARS-CoV-2 RNA was absent, and therefore the specimen is negative for COVID-19. In contrast, observing SARS-CoV-2 signal only indicates that SARS-COV-2 RNA was so abundant in the specimen that it is above the quantifiable range. In a mixed chromatogram of spike-in and SARS-CoV-2, the abundance of SARS-CoV-2 RNA is determined by the relative contributions of SARS-CoV-2 and spike-in signal intensities. (**B**-**D**) Synthetic viral genomic RNA was added to RT-PCR reactions at 10, 100, 1000, and 5000 GCE. The same dilutions were subjected to qSanger testing and RT-qPCR. (**B**) RT-qPCR exhibits a linear estimate across the dilution series, consistent with previous results. (**C**) Across the same dilution series, the ratio (R) of genomic sequence to spike-in sequence scales with RNA added. (**D**) When the qPCR estimate of abundance is compared to qSanger estimates of abundance, they exhibit a strong linear relationship, indicating that qSanger performs as well as qPCR in estimates of viral RNA abundance.

To determine the limit of detection of qSanger COVID-19, we performed assays on dilutions of SARS-COV-2 RNA corresponding to 0 GCE, 10 GCE, 100 GCE, 1000 GCE, or 5000 GCE. Four replicates at each dilution were assayed by both qSanger and qPCR. As expected, all 4 replicates of 0 GCE were negative for COVID-19 by qPCR, and addition of 10 or more SARS-CoV-2 GCE exhibited a clear logarithmic decrease in qPCR cycle threshold (Ct) (Fig. 2B). Similarly, no SARS-CoV-2 sequence was apparent on Sanger chromatograms for the NTC condition. No spike-in sequence was qualitatively discernable at the 5000 GCE dilution. Mixed bases were obviously present for both the 10 GCE and 100 GCE conditions suggesting that the limit of detection is about 10 GCE SARS-CoV-2.

We developed custom bioinformatic analyses to extract the relative abundance of SARS-CoV-2 and spike-in amplified products from Sanger chromatograms and automate analysis of qSanger chromatograms (see Methods). Briefly, peak amplitudes were assigned to either the spike-in or SARS-CoV-2 sequence at each base position, and a linear regression analysis was performed to determine the qSanger ratio between SARS-CoV-2 and spike-in trace intensities. qSanger ratios near 0 were recovered in the samples with 0 GCE, indicating the complete absence of SARS-CoV-2 RNA. (Fig. 1C). Since all of the SARS-CoV-2 RNA at 10 GCE or qSanger ratios of 3% or greater, this provided further evidence that the limit of detection of qSanger COVID-19 is ~10 GCE or fewer. Quantitative analysis of chromatogram peak heights was able to recover a qSanger ratio for even the 5000 GCE condition, and the qSanger ratios had excellent linearity over 10-5000 GCE (Fig. 2C). Furthermore, qSanger ratios were in good agreement qPCR Ct values (Fig. 2D).

### qSanger detects SARS-CoV-2 RNA when amplified directly from viral particles in transport medium without RNA purification

Since a major limitation for increasing testing capacity has been supply chain and lab workflow bottlenecks related to RNA extraction, we next attempted to detect SARS-CoV-2 directly from the specimen matrix (viral transport medium). There has been a previous report that RT-qPCR can be successfully performed when up to 3-7ul (total reaction volume of 20ul) of VTM without extraction is used as the template for RT-PCR [9]. We hypothesized that qSanger could be a more reliable method for detection of COVID-19 without RNA extraction because of *i*. increased robustness against PCR inhibitors in the specimen matrix since quantification of SARS-CoV-2 is performed via comparison with the spike-in internally control; *ii*. an improved limit of detection by adding more VTM to a correspondingly larger reaction size; and, *iii*. avoidance of false-positives by examining sequencing data for the correct spike-in or SARS-CoV-2 sequence. To test this hypothesis, we obtained reference materials in which either SARS-CoV-2 (positive control) or human RNA (negative control) is packaged inside of viral particles and suspended in VTM (Seracare). Since polymerase mixes can have varying resiliency to PCR inhibitors, we also evaluated both Luna RT-qPCR and OneTaq RT-PCR kits.

For both Seracare negative and positive control specimens, eight replicates each were performed on the cross-product of conditions for Luna vs OneTaq polymerases and direct VTM vs purified RNA, for 64 reactions total. 25ul of the Seracare specimen (corresponding to 125 GCE) was added to a 100ul total reaction volume. Additionally, 16 replicates of no-template controls were performed for each polymerase wherein nuclease free water was added to the reaction. All 32 NTC samples across all conditions were negative by qSanger assay and analysis (Fig. 3A). Nearly all Seracare negative control samples were determined to be negative by qSanger; indeterminate results were obtained from 2 purified RNA Luna specimens and 1 purified OneTaq specimen. All Seracare positive controls were identified by qSanger except for a OneTaq direct VTM specimen (Fig. 3A). Samples were classified as undetermined due to low chromatogram signal intensity (Signal to noise ratio < 500) or lack of sequence alignment. Possible reasons for undetermined chromatograms could be failure in RNA extraction, PCR amplification, or cycle sequencing. Since the majority of undetermined specimens used purified RNA, the possibility that the majority of assay failures were due to the RNA extraction process itself, perhaps by carryover of high salt buffers, should be considered.

**Figure 3.**
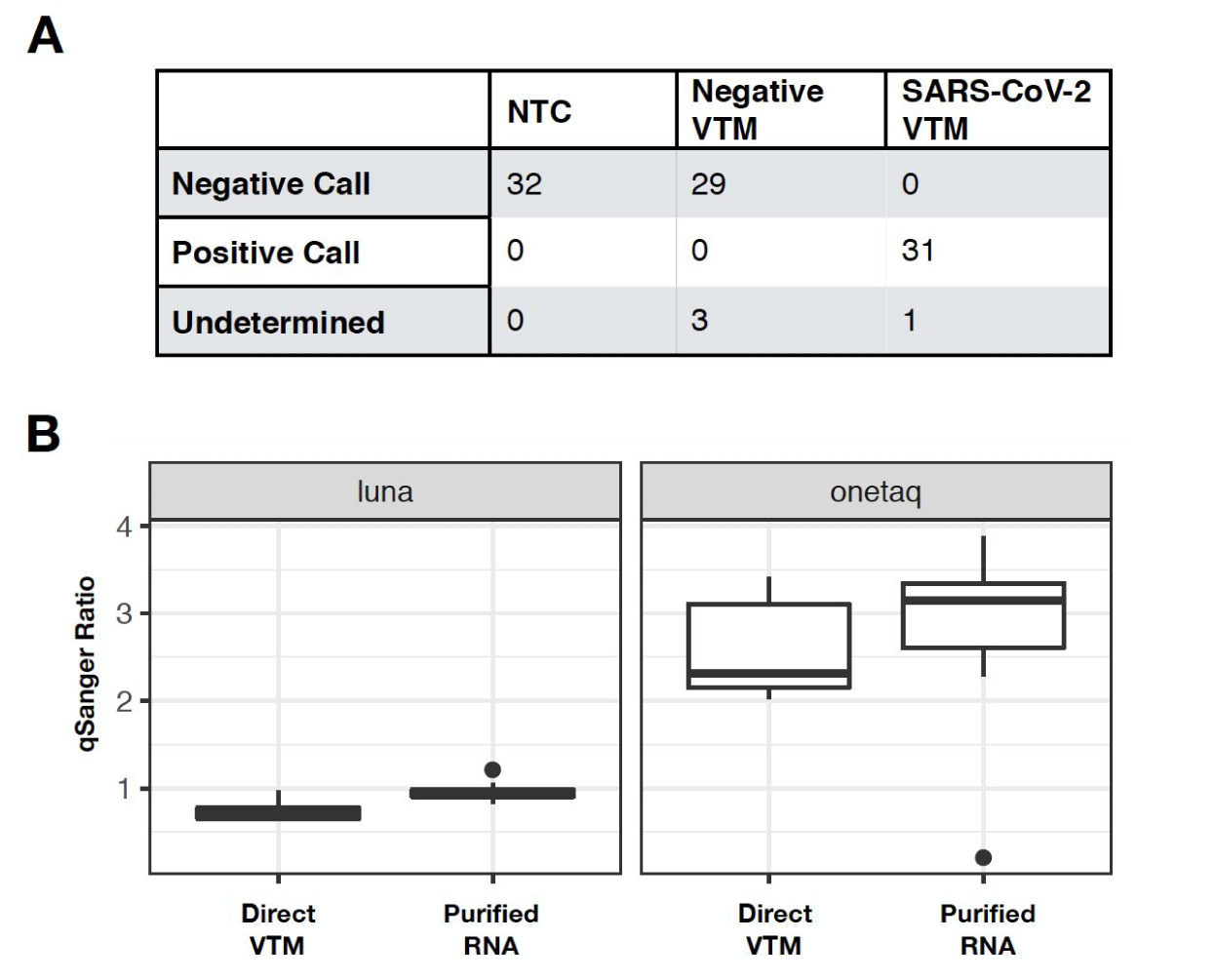
qSanger detects SARS-COV-2 RNA when amplified directly from viral particles in transport medium. (**A**) A total of 32 no-template controls, 32 negative control samples (Seracare) and 32 positive samples (Seracare) were assayed. All results were concordant with Seracare and NTC. Three samples were no-calls (undetermined) due to low signal-to-noise ratio in the sequencing results. (**B**) Positive Seracare samples were added to RT-PCR master mix either directly from the VTM or after purification with RNA extraction kit at 125 GCE. The ratio of reference and spike-in intensities were measured by custom data analysis. The mean qSanger ratio was 0.745 (± 0.043 s.e.m., n=8) for direct addition, and 0.97 (± 0.041 s.e.m., n=8) for purified. The coefficient of variation (CV) of positive seracare samples were measured for both Luna and OneTaq polymerase mixes. The CV for Luna direct VTM was 16.4% (n=8), and for Luna purified was 12.1% (n=8). This is consistent with the theoretical counting noise associated with quantifying ~100 molecules.

To further evaluate the feasibility of a direct VTM, extraction-free method for Sars-CoV-2 detection, we also examined the ability of qSanger to quantify the amount of viral particles in the Seracare positive control specimens (Fig. 3B). Since each reaction contained 125 GCE of Sars-CoV-2 and 200 GCE of spike-in, we expected to measure a qSanger ratio of 0.625. The OneTaq results yielded a qSanger ratio consistently around 2-3.5. This discrepancy could be due to a slightly more efficient amplification of the SARS-CoV-2 sequence compared to spike-in sequence. Remarkably, Luna polymerase mix yielded a qSanger ratio of 0.74 +/− 0.04 s.e.m. for VTM and a qSanger ratio of 0.97 +/− 0.04 s.e.m. for purified RNA, which is very close to the expected ratio. The ~20% difference is on par with typical imprecisions for pipetting and DNA quantification. The coefficient of variation (CV) for the VTM and RNA purified Luna assays were 12% and 16%, respectively. Notably, this is in good agreement with the imprecision associated with measuring ~125 molecules at the Poisson limit. Since Luna exhibited better accuracy and precision compared to OneTaq, and the Luna direct VTM method resulted in correct calls for all NTC, Seracare positive, and negative specimens without any failed reactions, subsequent experiments were performed with Luna polymerase mix.

Finally, we sought to demonstrate that omitting RNA extraction does not adversely affect qSanger sensitivity. We added 20 GCE (corresponding to 2x the LOD in Fig. 1) of viral particles in VTM containing either negative control or SARS-CoV-2 RNA (Fig. 4). Sanger chromatograms clearly showed the absence and presence of SARS-CoV-2 signal in the negative and positive controls, respectively (Fig. 4A). The qSanger assay correctly identified the negative and positive samples in 39 out of the 40 samples tested, with 1 negative control specimen returning an undetermined result due to sequencing failure (Fig. 4B). Overall, the excellent performance shown by our qSanger results on unextracted VTM vs. purified RNA with respect to absolute quantification accuracy, Poisson-limited coefficient of variation, and limit of detection that is comparable to gold-standard RT-qPCR, suggests that qSanger can be performed on unprocessed specimen matrix without any loss in performance. In fact, it might be possible that RNA-extraction free methods are more reliable because it eliminates the carryover risk of high-salt, PCR-inhibiting buffers used in RNA extraction procedures.

**Figure 4.**
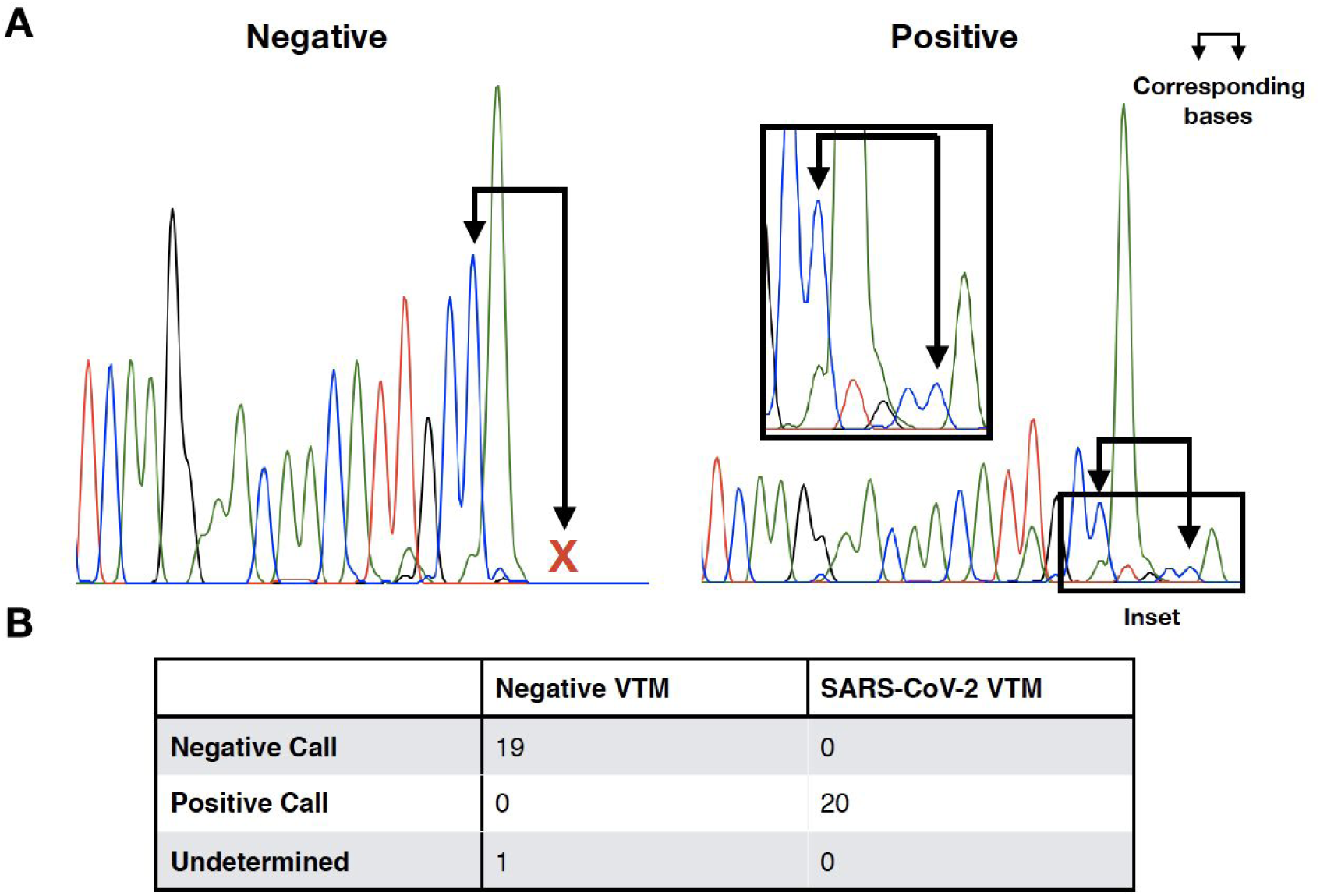
qSanger detects as little as 20 GCE without RNA purification. (**A**) Representative Sanger sequencing traces of negative control virus (left) and SARS-CoV-2 sequence containing virus (right). Even when the signal is too low to detect mixed bases, the 3’ offset caused by the deletion in the spike-in compared to the genomic sequence identifies positive samples. The sequencing peaks identified in the inset correspond to spike-in sequence offset by 4 bp (see paired arrows). (B) Twenty samples each of negative control virus and SARS-CoV-2 sequence containing virus were directly added to RT-PCR master mix with 100 spike-in molecules and Sanger sequencing was performed. All samples that successfully sequenced were accurately identified (one negative sample was undetermined due to sequencing failure).

## Discussion

The COVID-19 pandemic is one of the most deadly and disruptive public health emergencies that we have had to face in the last hundred years. To curb its rapid spread, most countries issued shelter-in-place rules which were able to flatten the pandemic curve, but at a significant economic cost [10]. Social distancing interventions can only be relaxed if a vaccine is developed, which is estimated to take 12 months to develop and widely distribute, or after widespread and rapid testing can be performed on symptomatic patients as well as anyone who has been identified as likely to have come into contact with those who test positive [11]. This suggests that for effective contact tracing, up to 50 additional people may need to be tested for each positive case. However, on April 6, 2020, the US daily testing capacity was only ~100,000 to ~125,000 tests when 30,000 patients tested positive [12]. This suggests that US testing capacity might need to be increased by a factor of 10x or more for effective contact tracing. However, most COVID-19 tests that are currently available rely on variations on qPCR and thus suffer from the same supply chain issues. In particular, the limited availability of RNA extraction kits has been an important problem for increasing COVID-19 testing capacity [13].

Here, we describe a novel quantitative Sanger (qSanger) assay that can detect COVID-19 without RNA extraction. We show that qSanger performs as well as qPCR in estimates of viral RNA abundance and consistently detects as low as 10-20 viral genome copy equivalents, even when VTM is added directly into the reaction mix without RNA extraction. Because qSanger is an end-point PCR reaction with an internal spike-in control, it is more robust to inhibitors that can exist in VTM, and failures in amplification result in undetermined results that require a repeat reaction, as opposed to false negatives that would be obtained by qPCR. It also has higher specificity than qPCR, as the examination of the sequencing information can be used to distinguish similar viruses and rule out false-positives due to non-specific amplifications.

Since qSanger has an extremely high specificity enabled by the sequencing information, it can be used for routine testing of asymptomatic individuals with high positive predictive value (PPV). This can be a new paradigm of routine and repeated testing of individuals who are at high risk for contracting disease, e.g., hospital staff, or those who are older or with comorbidities. Early detection can improve individual healthcare outcomes and also enable relaxation of population scale non-pharmaceutical interventions like social distancing measures.

In addition not requiring RNA extraction kits, our qSanger-based COVID-19 assay has a number of additional advantages compared to existing qPCR-based tests for COVID-19. Importantly, qSanger thermal cycling occurs in higher-throughput end-point PCR instruments, rather than specialized qPCR instruments, and the sequencing can be run in automated Sanger sequencers with plate feeders such as Applied Biosystems 3730xl DNA Analyzers which have the capacity to sequence 3840 samples per day. The large existing install-base of end-point PCR and high-throughput Sanger instruments throughout the US and the world [14] supports rapid scale-up of qSanger-COVID-19 assays without requiring any new device or instrument manufacturing. Given that Sanger sequencing is still the most widely used method of clinical sequencing worldwide, the widespread adoption of qSanger-COVID-19 assay described here can create >1M COVID-19 testing capacity per day.

More broadly, qSanger can enable an even higher volume of population-scale testing if clinical laboratories and Sanger sequencing centers are allowed to collaborate for COVID-19 testing. While Sanger sequencers exist in all molecular diagnostic laboratories, they are most commonly utilized in high volume in genome centers, academic sequencing core facilities, and commercial Sanger sequencing service laboratories. In this model, clinical laboratories could buy 96-well master-mix reaction plates that simply require the addition of each patient sample into a reaction well in a BioSafety Cabinet and PCR thermocycling, whereas the sequencing service laboratory would sequence the samples for next-day results. This would enable a rapidly deployed and distributed population-scale testing.

qSanger also has a number of other advantages that may prove to be invaluable as we learn more information about SARS-CoV-2. Since SARS-CoV-2 mutates quickly, the availability of sequence information can be used to identify growing clusters of mutations and aid with contact tracing via phylogenetic analysis of the mutation data. Longer sequences can be designed to capture a wide range of mutations in the qSanger reaction, as the virus mutates and creates sub-strains with different clinical implications.

Moreover, as opposed to relative measurements obtained by qPCR, absolute measurements of viral load are obtained in qSanger due to the known molecular count of the spike-in. Quantification of viral abundance in a sample may prove to be useful for determining who is infectious, as well as for more accurate environmental monitoring. The quantitative dynamic range of qSanger can be broadened from 0-5000 GCE to 10-2,500,000 GCE by employing two qSanger reactions with different molecular levels of spike-ins.

At the time of the Human Genome Project, Sanger instruments were the automated state-of-the-art devices that generated more sequencing information in a single day than entire laboratories could produce in a year [14]. They accelerated the release of the first human genome by five years and ushered in an era with exponentially decreasing sequencing costs [14–15]. With the qSanger method presented here, they can be repurposed to enable the millions of COVID-19 diagnostics needed to suppress this global pandemic.

## Methods

### Primer and spike-in

Spike-in sequences were designed using the viral genomic region corresponding to the CDC designed N3 qPCR assay. Spike-in molecules have sequence identical to SARS-CoV-2 sequence (LC528232) including base positions 28216 to 29280 but lacking bases 28715-28718. Primers flanking the deletion were used for amplification. Sequencing was performed using a primer containing the forward amplification binding region.

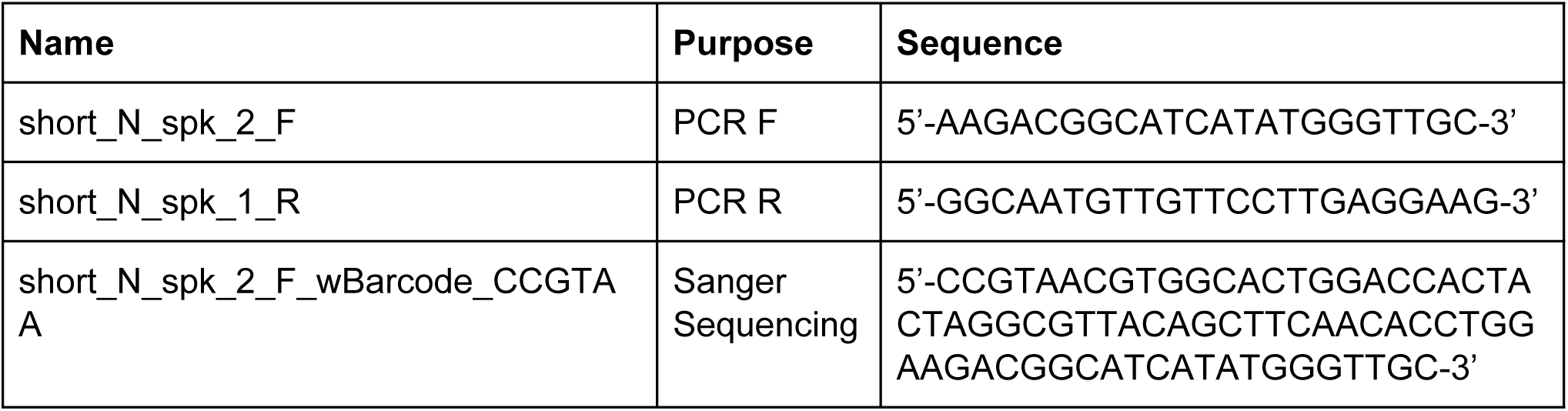

#### Samples

Synthetic SARS-CoV-2 genomic RNA from Twist Biosciences was used for RNA detection linearity experiments. AccuPlex SARS-CoV-2 Reference Material Kit manufactured by Seracare (cat. # 0505-0126) was used as a proxy for clinical samples.

#### Viral Purification

Viral purification was performed using PureLink Viral RNA/DNA Mini Kit from ThermoFisher Scientific using 500 µL (at 5000 GCE/mL) of Accuplex Positive or Negative sample. GCE input estimates from purified RNA were estimated by corresponding fraction of eluent assuming 100% recovery.

#### qPCR

qPCR were performed using the N1 primers and TaqMan probes provided in the 2019-nCoV RUO Kit manufactured by IDT (cat. # 10006605). Amplification was performed as described in the CDC EUA protocol. Briefly, 2 µL of synthetic RNA template diluted in RNAse-free TE + 0.05% Tween-20 to 5, 50, 500, or 2500 GCE/µL were added to each reaction.. RNA samples were combined with water, TaqPath 1-Step RT-qPCR Master Mix, primers (1.5 µL to a final concentration of 500 nM), and probes to a total final volume of 20 µL. The reaction mixture was amplified and probe fluorescence was detected using a Mastercycler ep realplex Real-time PCR System. The first cycle above threshold was estimated (Ct) was performed with default settings using *realplex* software.

#### qSanger Amplification

Reverse transcription and amplification for figure 2 was performed using OneTaq One-Step RT-PCR Kit from NEB (cat. # E5315S). Both the OneTaq One-Step RT-PCR Kit and Luna Universal One-step RT-qPCR Kit (cat. # E3005E) were used for figure 3. Figure 4 was performed exclusively with the Luna Universal One-step RT-qPCR Kit. Buffer and enzyme were used according to manufacturer recommendations for 100 µL total volume. All reactions contained Tween-20 at a final concentration of 1% v/v, 500 nM final concentration of each amplification primer, and 100 GCE of synthetic dsDNA spike-in molecules. Synthetic RNA samples were added at 2 µL/reaction and synthetic sample was added to achieve the appropriate number of viral particles. Thermocycling was performed using an Applied Biosystems Veriti Thermal Cycler with the following cycling programs:

| # of Cycles | OneTaq |  | Luna |  |
| --- | --- | --- | --- | --- |
| 1x | 48°C | 20:00 | 55°C | 20:00 |
| 1x | 94°C | 1:00 | 95°C | 1:00 |
| 40x | 94°C | 0:20 | 95°C | 0:20 |
|  | 55°C | 1:00 | 55°C | 1:00 |
|  | 68°C | 1:00 | 60°C | 1:00 |
| 1x | 68°C | 5:00 | 68°C | 5:00 |

#### Sanger sequencing

Sanger sequencing was performed by Sequetech Corporation using the BigDye Terminator Cycle Sequencing Kit and capillary electrophoresis was performed using a Applied Biosystems 3730xl DNA Analyzer.

#### Data analysis

For concordance calls, Sanger sequencing was analyzed using the following procedure. The primary base sequence based on automatic calling were aligned to the viral genome. If the aligned sequence matched the viral genomic sequence without any deletion, then the sample is called positive for viral RNA. If the sequence does not match the reference, then the signal to noise ratio is checked with any less than 500 indicating insufficient signal which returns an indeterminate result. If signal to noise is greater than 500, then the ratio of genomic sequence to spike-in sequence is quantified by performing robust linear regression of genomic peak heights to spike-in peak heights.

Quantitation for figure 2 was performed with the same quality check as above but quantifying the terminal 6 bases of reference and spike-in sequence for all primary sequences, regardless of whether genomic, spike-in, or mixed sequence dominates. All analysis was performed using custom scripts in R, employing the *seqinr* and *tidyverse* packages.

## Author Contributions

D.S.T. conceived the project, D.C.-B, O.A., and D.S.T. designed the experiments. A.M.B. and D.C.-B performed the experiments. D.C.-B. and D.S.T. analyzed the data. D.C.-B., O.A. and D.S.T. wrote the manuscript.

## Acknowledgements

We thank Paul Buchheit and Y Combinator for their financial support of the project, employees of Genewiz, Sequetech, IDT for support with and prioritization of our orders, Shan Riku and Oscar Cabot for their help with the distribution of the manuscript.

